# Structural Basis of Stu2 Recruitment to Yeast Kinetochores

**DOI:** 10.1101/2020.10.31.362574

**Authors:** Jacob A. Zahm, Michael G. Stewart, Joseph S. Carrier, Stephen C. Harrison, Matthew P. Miller

**Author notes:** Theses authors contributed equally to this work (Jacob A. Zahm, Michael G. Stewart). Correspondence (S.C.H.), (M.P.M.).

## Abstract

Accurate chromosome segregation during cell division requires engagement of the kinetochores of sister chromatids with microtubules emanating from opposite poles of the mitotic spindle. In yeast, these “bioriented” metaphase sister chromatids experience tension as the corresponding microtubules (one per sister chromatid) shorten. Spindle-assembly checkpoint signaling appears to cease from a kinetochore under tension, which also stabilizes kinetochore-microtubule attachment in single-kinetochore experiments in vitro. The microtubule polymerase, Stu2, the yeast member of the XMAP215/ch-TOG protein family, associates with kinetochores in cells and contributes to tension-dependent stabilization, both in vitro and in vivo. We show here that a C-terminal segment of Stu2 binds the four-way junction of the Ndc80 complex (Ndc80c) and that amino-acid residues conserved both in yeast Stu2 orthologs and in their metazoan counterparts make specific contacts with Ndc80 and Spc24. Mutations that perturb this interaction prevent association of Stu2 with kinetochores, impair cell viability, produce biorientation defects, and delay cell-cycle progression. Ectopic tethering of the mutant Stu2 species to the Ndc80c junction restores wild-type function. These findings show that the role of Stu2 in tension sensing depends on its association with kinetochores by binding with Ndc80c.

## INTRODUCTION

Equal partitioning of duplicated chromosomes during cell division preserves integrity of the genome in each of the two daughter cells. “Bioriented attachment” of sister chromatids to opposite poles of the mitotic spindle in turn ensures that when a cell enters anaphase, each pair of sister chromatids segregates accurately (reviewed in Cheeseman, 2014). When correctly bioriented, a pair of cohesin-linked sister chromatids will be under tension, from forces exerted by the shortening of opposing microtubules and transmitted through the kinetochores that connect spindle microtubules to chromosome centromeres. “Tension-sensing” and correction of erroneous attachments are thus critical mediators of genome integrity.

The Aurora B kinase (Ipl1 in budding yeast) is the immediate agent of error correction (Biggins et al., 1999; Cheeseman et al., 2002; DeLuca et al., 2006; Hauf et al., 2003; Tanaka et al., 2002). In the absence of tension, Ipl1 phosphorylates Ndc80 (DeLuca et al., 2006; Hauf et al., 2003), an essential part of the microtubule-contacting apparatus of a yeast kinetochore, and several other kinetochore substrates (Biggins and Murray, 2001; Biggins et al., 1999; Cheeseman et al., 2002). Phosphorylation induces dissociation of a kinetochore from its associated microtubule, interrupting the incorrect attachment and allowing the attachment search to “try again”. Ndc80 is part of a rod-like assembly, the Ndc80 complex, a heterotetramer composed of two coiled-coil heterodimers, joined end-to-end in parallel orientations (Alushin et al., 2010; Ciferri et al., 2008; Wei et al., 2005; Wilson-Kubalek et al., 2008). The globular domain at the N-terminal end of Ndc80:Nuf2 contacts microtubules; the globular domain at the C-terminal end of Spc24:Spc25 interacts, through two distinct adaptors, with the chromatin-proximal inner kinetochore. The Ipl1 substrate residues of Ndc80 are in an N-terminal extension that contributes, along with a globular, calponin homology domain, to the microtubule interface (DeLuca et al., 2006; Umbreit et al., 2012).

A further component of the response to erroneous attachment is the plus-end microtubule polymerase, Stu2 (the yeast ortholog of XMAP215/ch-TOG in metazoans). Its polymerization activity depends on plus-end binding, through a set of N-terminal TOG domains (Fig. 1A) (Al-Bassam et al., 2006; Ayaz et al., 2012, 2014; Fox et al., 2014), but its kinetochore association is independent of microtubules (Hsu and Toda, 2011; Kakui et al., 2013; Miller et al., 2016, 2019; Vasileva et al., 2017). In single-molecule experiments *in vitro*, kinetochores under low tension detach more frequently from microtubules than under higher tension (Akiyoshi et al., 2010), but only when Stu2 is present (Miller et al., 2016). Cells lacking Stu2 have error-correction defects, including detached kinetochores, loss of biorientation, and spindle-assembly checkpoint-dependent cell-cycle delay, although previous work has been unable to separate these phenotypes from pleiotropic effects arising from loss of Stu2 microtubule polymerase activity (Al-Bassam et al., 2006; Humphrey et al., 2018; Kosco et al., 2001; Miller et al., 2016; Pearson et al., 2003; Severin et al., 2001; Wang and Huffaker, 1997).

**Figure 1.**
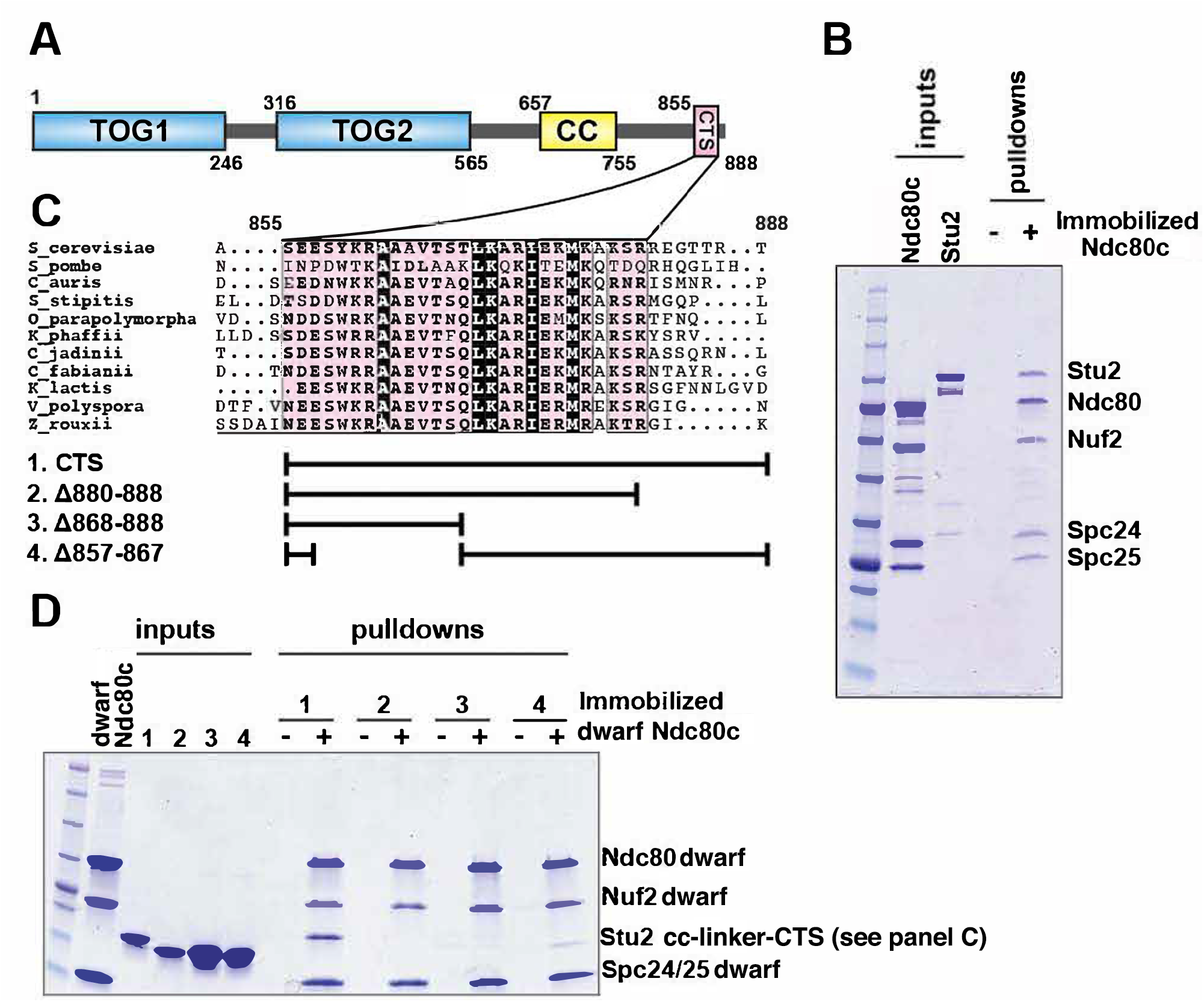
Binding of Stu2 CTS with Ndc80c. **A.** Schematic representation of the domain structure of Stu2, showing the tubulin-binding TOG domains, the dimeric coiled-coil, and the C-terminal segment (CTS). Molecular mass markers: 245,190,135,100,80,58,46,32,25,22,17,11kDa from top to bottom (New England Biolabs). **B.** Association of full-length Ndc80c and Stu2. Ndc80c was immobilized on Co-NTA agarose and incubated with Stu2. After extensive washing, bound proteins were eluted with buffer containing 400 mM imidazole. **C.** Multiple sequence alignment showing conservation of the CTS among budding and fission yeasts. The bars below the alignment correspond to the *Saccharomyces cerevisiae* sequence in the alignment and show the constructs used in the pulldown experiments in **D**. The blank parts of each line represent deletions. **D.** Binding of dwarf Ndc80c andStu2 CTS. The Stu2 constructs used in this experiment consist of the Stu2 coiled-coil domain, followed by a glycine-serine linker, followed by the regions of the CTS indicated by the bars in **C.** Dwarf Ndc80c was immobilized on Co-NTA agarose and incubated with Stu2. After extensive washing, bound proteins were eluted with buffer containing 400 mM imidazole.

Stu2 associates with Ndc80c (Miller et al., 2016, 2019). We describe experiments here in which we found that a C-terminal segment (CTS) of Stu2 bound specifically at the junction of the Ndc80:Nuf2 and Spc24:Spc25 heterodimers and that conserved features of the four-chain overlap at that junction were important for the interaction. Mutations in Stu2 that disrupted this interaction blocked association of Stu2 with isolated kinetochores *in vitro*, impaired cell viability, and caused defects in chromosome biorientation and in timely cell-cycle progression. Ectopic tethering of the mutant Stu2 to Ndc80c rescued these defects, allowing us to assign them to the interaction with Ndc80 rather than to interactions with other binding partners, such as Bik1 and Spc72 (Usui et al., 2003; Wolyniak et al., 2006). We conclude that Stu2 stably associates with the Ndc80c four-way junction and that this association is critical for establishing bioriented kinetochore attachments and for maintaining cell viability. These findings are consistent with the notion that kinetochore-associated Stu2 is a central component of the tension-sensing mechanism in budding yeast.

## RESULTS

### Interaction of a conserved Stu2 C-terminal segment with Ndc80c

To specify interactions that stabilize a Stu2:Ndc80c complex, we determined which regions of Stu2 are both necessary and sufficient for binding to Ndc80c (Fig. 1A). Initial pulldown experiments showed that recombinant full-length Stu2 bound directly to immobilized recombinant full-length Ndc80c (Fig. 1B). Both the flexibility and elongated shape of Ndc80c would hinder structural studies of the full-length proteins. We instead used the “dwarf” version of Ndc80c in which the coiled-coil shaft between the globular heads has been shortened to facilitate crystallization (Valverde et al., 2016). In pulldown experiments, we found that Stu2 binds dwarf Ndc80c as efficiently as it does full-length Ndc80c (Fig 1B). Subsequent pulldown experiments used immobilized dwarf Ndc80c and a construct comprising the Stu2 dimeric coiled-coil domain followed by a glycine-serine linker, joined in turn to the Stu2 C-terminal segment or fragments of it (Humphrey et al., 2018; Miller et al., 2019; Usui et al., 2003; Wang and Huffaker, 1997). We found that a segment at the C-terminus of Stu2, most of which is conserved among budding and fission yeast and referred to here as the “C-terminal segment” (CTS) (Fig. 1A), is required for binding Ndc80c. Deletion of either an N-terminal portion of the CTS (residues 857-867), a C-terminal portion of the CTS (residues 868-888), or the stretch following the CTS that lacks conservation (residues 880-888) resulted in loss of *in vitro* binding (Fig. 1C,D). These data show that a dimeric construct bearing the C-terminal 33 residues of Stu2 is sufficient for Ndc80c binding.

### Structure of Stu2 in complex with dwarf Ndc80c

We co-crystallized the dwarf Ndc80c with a 33-mer Stu2 peptide (residues 855-888) and determined the structure of the complex by molecular replacement using the published dwarf structure as a search model (Fig. 2A). Clearly defined density in the resulting map corresponded to residues 862-888; there was no density corresponding to the N-terminal 7 residues (Fig. 2B), and the C-terminal residues, 881-888, were part of a crystal-packing contact with the Nuf2 globular head of an Ndc80c symmetry mate (Fig. 2B). The peptide features in the map were weaker than those for the Ndc80c, probably owing to its relatively modest affinity. To resolve a potential ambiguity in the sequence register of the model, we determined the structure of a complex with a peptide containing selenomethionine in place of the methionine at position 876. A strong anomalous difference peak confirmed that we had chosen the register correctly (Fig. 2B).

**Figure 2.**
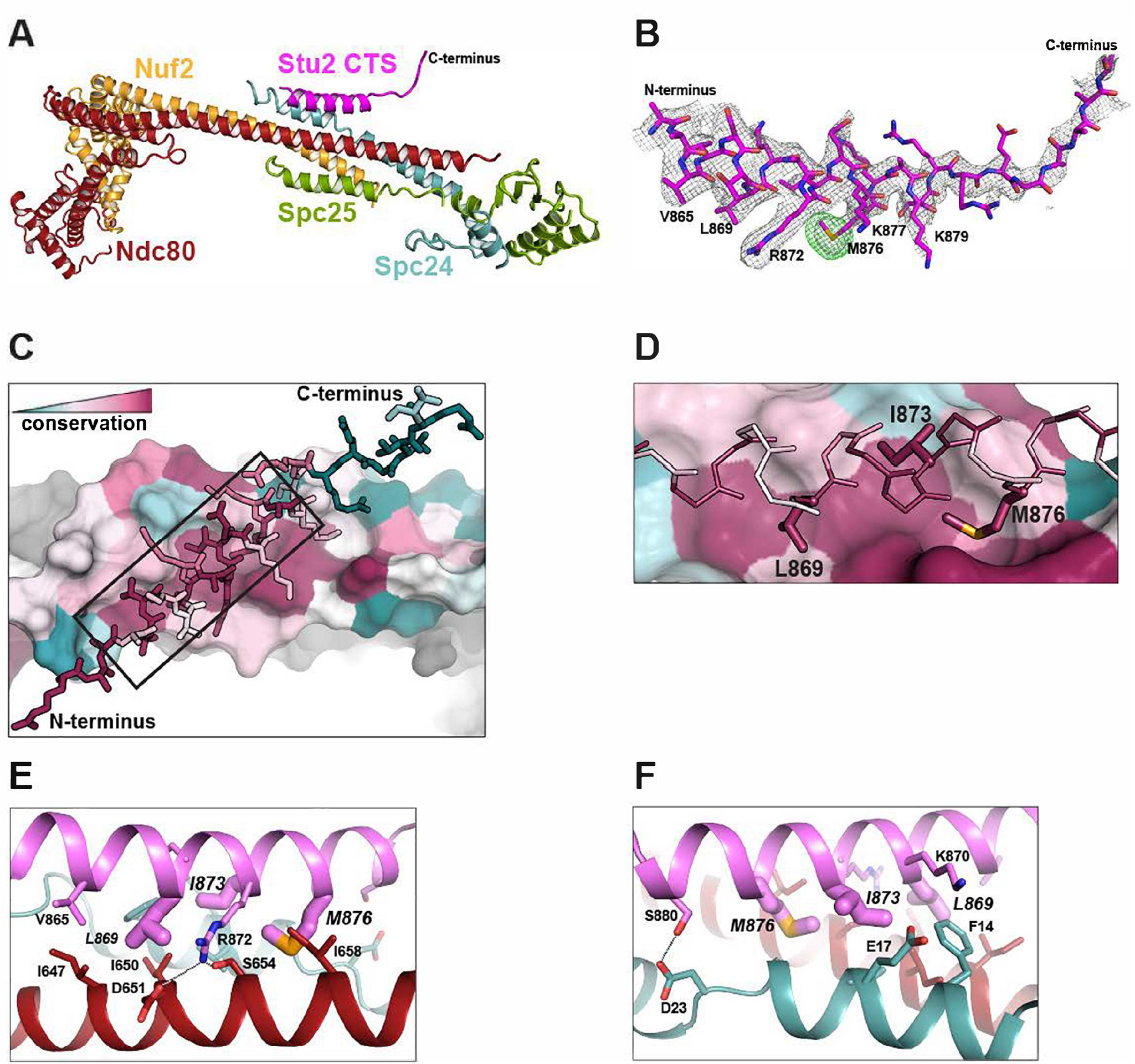
Structure of the Stu2 CTS bound to the dwarf Ndc80c. **A.** The Stu2 CTS (residues 855 – 888) bound to the dwarf Ndc80c. The peptide binds at the 4-way junction of Ndc80, Nuf2, Spc24 and Spc25 in a groove between Ndc80 and Spc24. **B.** Model of the Stu2 peptide built into the 2Fo-Fc map (grey mesh); anomalous difference map (green mesh), contoured at 8σ, showing the postion of SeMet in the peptide. **C.** Conservation of residues at the contacts of Stu2 CTS, Ncd80, and Spc24, shaded from red (conserved) to blue (variable). Nuf2 and Spc25 are in gray. Ndc80c components in surface representation; Su2 in stick representation. The N and C-termini of the Stu2 peptide are indicated. The expanded region depicted in **D** is shown as a black box. **D.** Expanded view of the Ndc80-Spc24 surface corresponding to the boxed region in panel C, showing conserved pockets for the three hydrophobic residues of Stu2 discussed in the text. Coloring as in **C**. **E-F.** Detailed views of the contacts between the Stu2 peptide and Ndc80 and Spc24.

The peptide binds at the junction of the two heterodimeric subcomplexes, contributing a fifth a helix, and packs at about 30° to the main axis of the four overlapping helices from Ndc80, Nuf2, Spc24 and Spc25. Four conserved hydrophobic-residue side chains in the Stu2 CTS (V865, L869, I873 and M876) fit into hydrophobic pockets lined by conserved residues of Ndc80 and Spc24 (Fig. 2A-F), and in a conserved polar contact, the side chain of Stu2 R872 inserts between D651 and S654 on Ndc80 (Fig. 2E).

To confirm the correspondence of the Stu2:Ndc80c interface in the crystal structure with binding in solution, we made point mutations in Stu2 and carried out pulldowns with immobilized dwarf Ndc80c, using the same Stu2 constructs described above, except for the point mutations. For the binding experiments we mutated both M876 and I873 to alanine. The immobilized dwarf Ndc80c did not bind either mutant version of the Stu2 C-terminus (Fig. 3A). These experiments show that the details of interaction between Stu2 and Ndc80c in the crystal lattice occur in solution as well.

**Fig 3.**
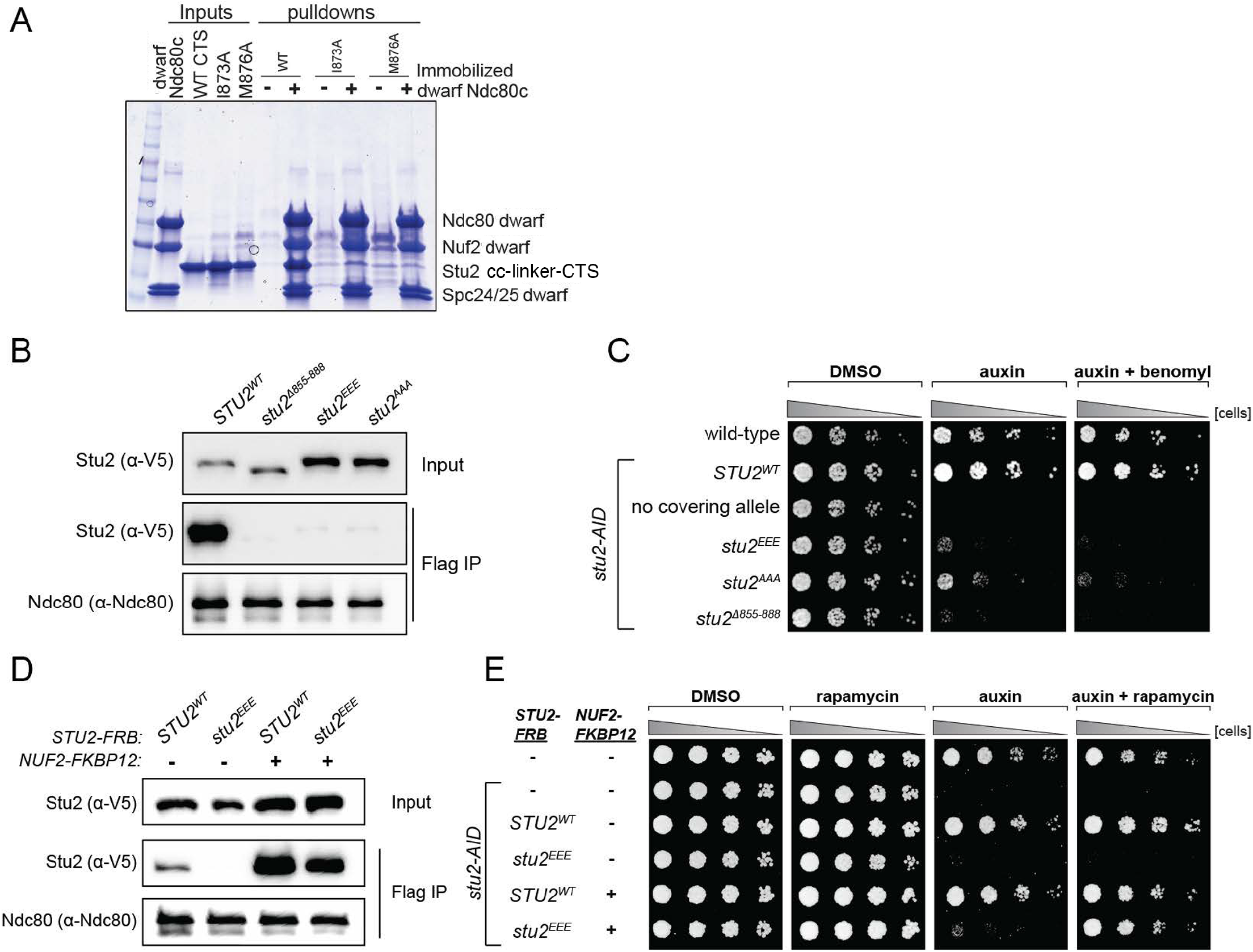
Effects of Stu2 mutations at the interface with Ndc80c on kinetochore association and cell viability and their rescue by re-tethering. **A.** Effect of mutations at the binding interface. The Stu2 constructs contain the Stu2 coiled-coil domain, a glycine-serine linker, and either native or mutated CTS. Dwarf Ndc80c was immobilized on Co-NTA agarose, incubated with Stu2, washed, and eluted with 400 mM imidazole. **B.** Exponentially growing *stu2-AID* cultures expressing an ectopic copy of *STU2* (*STU2^WT^*, M622; *stu2^Δ855-888^*, M653; *stu2^L869E,I873E,M876E^* or *stu2^EEE^*, M1444; *stu2^L869A,I873A,M876A^* or *stu2^AAA^*, M1576) and also expressing from the genomic locus Dsn1-6His-3Flag were treated with auxin 30 min prior to harvesting. Kinetochore particles were purified from lysates by anti-Flag immunoprecipitation (IP) and analyzed by immunoblotting. **C.** Wild-type (M3), *stu2-AID* (no covering allele, M619), and *stu2-AID* cells expressing various *STU2-3V5* alleles from an ectopic locus (*STU2^WT^*, M622; *stu2^EEE^*, M1444; *stu2^AAA^*, M1576; *stu2^Δ855-888^*, M653) were serially diluted (5-fold) and spotted on plates containing DMSO, 500 μM auxin, or 500 μM auxin + 5 μg/mL benomyl. **D.** Exponentially growing *stu2-AID fpr1Δ TOR1-1* cultures expressing an ectopic copy of *STU2-FRB* with wild-type *NUF2* (*STU2^WT^*, M1513; *stu2^EEE^*, M1515) or with *NUF2-FKBP12* (*STU2^WT^*, M1505; *stu2^EEE^*, M1507) were treated with rapamycin and auxin 30 min prior to harvesting. Protein lysates prepared, subjected to a-Flag IP, and analyzed as in **B**. **E.** *fpr1Δ TOR1-1* cells (M1375), and *stu2-AID fpr1Δ TOR1-1* cells (no covering allele, M1476), and *stu2-AID fpr1Δ TOR1-1* cells expressing *STU2-FRB* alleles at an ectopic locus with and without *NUF2-FKBP12* (M1513; M1515; M1505; M1507 – same genotypes as in **D**) were serially diluted (5-fold) and spotted on benomyl plates containing DMSO, 50 ng/mL rapamycin, 500 μM auxin, or 500 μM auxin + 50 ng/mL rapamycin.

### Kinetochore-associated Stu2 is required for cell viability

We generated strains containing point mutations at two positions tested *in vitro* and also at position L869, positions of three of the four conserved hydrophobic residues on Stu2 that contact Ndc80 and Spc24. For *in vivo* assays, we engineered an auxin-inducible degron in which addition of auxin would induce degradation of endogenous Stu2-AID, uncovering the phenotype of an ectopic mutant allele (Miller et al., 2019). We assessed in these strains the effects of the several Stu2 mutations on Stu2-kinetochore association as well as on cell viability. Kinetochore co-immunoprecipitations from *stu2-AID* cells harboring either hydrophobic-to-charged or alanine mutations of these residues, *stu2^L869E,I873E,M876E^* (*stu2^EEE^*) or *stu2^L869A,I873A,M876A^* or (*stu2^AAA^*), showed loss of Stu2-kinetochore binding. These same mutations resulted in a severe cell viability defect in the presence of auxin, exacerbated by addition of low concentrations of the microtubule destabilizing drug benomyl (Fig. 3B-C). The individual *stu2* mutations also all had reduced cell viability (Fig. S3A-B). Thus, the hydrophobic residues in the CTS of Stu2, which contact Ndc80c in the crystal structure, are required for Stu2-kinetochore binding and cell viability *in vivo*.

The observed growth defect resulting from disruption of Stu2-kinetochore binding could, in principle, be due to loss of microtubule polymerase activity or to association with other Stu2 CTS binding proteins (Gunzelmann et al., 2018; Usui et al., 2003; Wolyniak et al., 2006). To determine whether the viability defect of the most penetrant mutant, *stu2^EEE^*, was due solely to loss of kinetochore binding or whether other Stu2 activities were involved, we tested rescue of the kinetochore binding of *stu2^EEE^* by rapamycin in cells expressing suitably engineered FRB and FKBP12 fusion proteins. We generated strains containing *stu2-AID* with *NUF2-FKBP12* at the endogenous *NUF2* locus and *STU2-FRB* alleles at an ectopic locus (Fig S4). We chose Nuf2 as the target for FKBP12 fusion because of the proximity of the C-termini of Nuf2 and Stu2 in the crystal structure. The cells also contained a mutant *TOR1* allele (*TOR1-1*) and *fpr1D*, to avoid inhibiting cell growth by the rapamycin treatment (Haruki et al., 2008). Cells expressing *stu2^EEE^-FRB* were deficient for Stu2-kinetochore binding as expected. Expressing *NUF2-FKBP12* and adding rapamycin in culture rescued both the Stu2 binding defect (Fig. 3D) and the cell viability defect of *stu2^EEE^* (Fig. 3E). Because rapamycin addition rescued both defects, we infer that loss of kinetochore localization is the cause of viability loss. Thus, *stu2^EEE^* is a true separation of function mutation that allows us to assess the cellular consequences of disrupting Stu2-kinetochore association.

### Proper chromosome biorientation and timely cell cycle progression require kinetochore-associated Stu2

The cellular phenotypes of the *stu2^EEE^* mutant are consistent with a role for kinetochore-associated Stu2 in error correction. If so, cells defective in Stu2-kinetochore binding should also have defective chromosome biorientation. We assessed chromosome biorientation by using a methionine-repressible *CDC20* allele (*pMET-CDC20*) to arrest cells in metaphase that also carry a fluorescently marked centromere of chromosome III (Straight et al., 1996). Under these conditions, opposing spindle pulling forces cause bioriented sister chromatids to separate, appearing as two distinct GFP puncta (Pearson et al., 2001). The frequency of biorientation in *STU2^WT^* cells was at the level usually found in this assay (44±2%, Fig 4A-B), but it was significantly lower in *stu2^EEE^* cells (16±1%, p = 0.0002).

**Fig 4.**
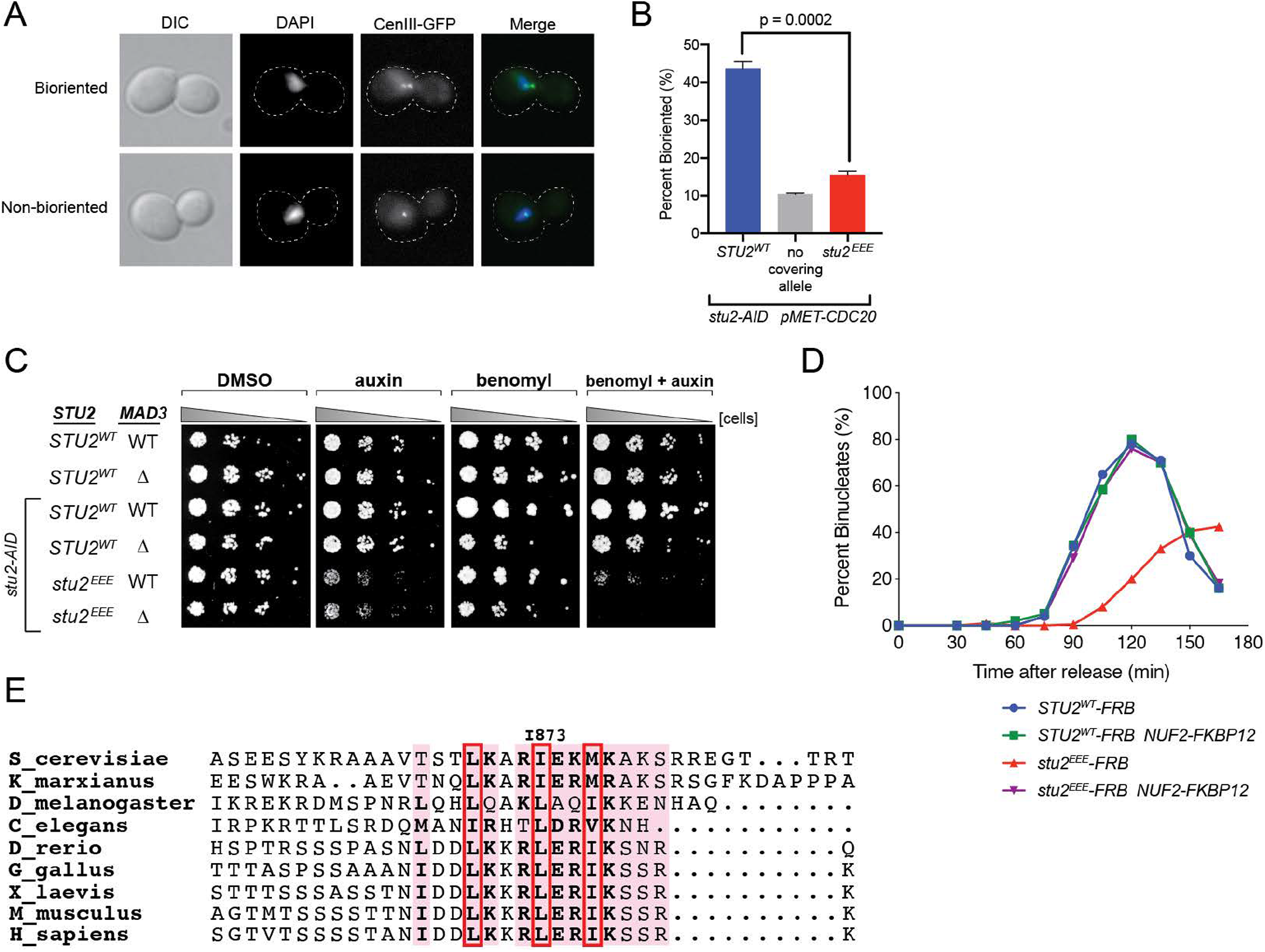
Cellular phenotypes of *stu2^EEE^* and conservation of key residues in multicellular eukaryotes. **A.** Exponentially growing *stu2-AID pMET-CDC20* cultures with an ectopically expressed *STU2* allele (*STU2^WT^*, M1154; *stu2^EEE^*, M1610) or no ectopic allele (no covering allele, M1153) and also containing cenIII marked with GFP (*cenIII-lacO lacI-GFP*) were arrested in methionine + auxin containing media for 2.5 hours. Representative micrographs for bioriented and nonbioriented cells shown. **B.** Percent bioriented cells was measured for cultures described in **A**. 3 replicates of n=200 cells shown; p value determined with unpaired t-test. **C.** Wild-type cells (M3), cells with spindle checkpoint mutation (*mad3Δ*, M36), and *stu2-AID* cells expressing an ectopic copy of *STU2* without and with a spindle checkpoint mutation (*STU2^WT^*, M622; *STU2^WT^ mad3Δ*, M1622; *stu2^EEE^*, M1444; *stu2^EEE^ mad3Δ*, M1541) were serially diluted (5-fold) and spotted on plates containing DMSO, 500 μM auxin, 5 μg/mL benomyl, or 500 μM auxin + 5 μg/mL benomyl. **D.** *stu2-AID* cells expressing *STU2-FRB* alleles at an ectopic locus with *NUF2^WT^* or *NUF2-FKBP12* (*STU2^WT^-FRB*, M1513; *stu2^EEE^-FRB*, M1515; *STU2^WT^-FRB NUF2-FKBP12*, M1505; *stu2^EEE^-FRB NUF2-FKBP12*, M1507) were released from a G1 arrest into auxin and rapamycin containing media. Cell cycle progression determined by the accumulation of binucleate cells. **E.** Multiple sequence alignment generated using the Stu2 C-terminus and C-termini from Stu2 eukaryotic homologs. Residues are colored by percent conservation. Hydrophobic residues important for Ndc80c binding are boxed in red.

Cells defective in error correction depend on the spindle checkpoint to prevent chromosome missegregation and have a delayed metaphase-anaphase transition (Biggins and Murray, 2001; Shonn et al., 2003; Stern and Murray, 2001). We indeed found a significant synthetic growth defect of *stu2^EEE^* with deletion of spindle checkpoint component *MAD3* in the presence of low concentrations of benomyl, likely due to severe aneuploidy in *stu2^EEE^ mad3Δ* cells (Fig 4C). To examine the timing of cell cycle progression in these cells, we released them from a G1 arrest and monitored nuclear divisions. Consistent with error correction defects, cells expressing *stu2^EEE^-FRB* had greatly delayed anaphase onset, rescued by tethering *stu2^EEE^*-FRB to the kinetochore with *NUF2-FKBP12* and rapamycin (Fig 4D & S4E). We conclude that correction of erroneous kinetochore-microtubule attachments in mitosis depends on kinetochore-bound Stu2. Moreover, rescue of the defect by ectopic tethering shows that CTS binding has a replaceable localization function and that the effector function for error correction lies elsewhere in the Stu2 molecule.

## DISCUSSION

Previous studies have shown that Stu2, a microtubule plus-end polymerase, associates with kinetochores during mitosis in yeast cells (He et al., 2001; Hsu and Toda, 2011; Kakui et al., 2013; Miller et al., 2016, 2019; Vasileva et al., 2017) and contributes to the response to tension of kinetochores *in vitro* (Miller et al., 2016). The findings reported here indicate that kinetochore-bound Stu2 indeed contributes to error correction in response to the absence of tension *in vivo*. We determined the structure of a C-terminal peptide from Stu2 bound with a shortened construct of Ndc80c that preserves the junction between the two heterodimers it comprises (Valverde et al., 2016). We then used information from that structure to generate mutant yeast strains harboring *stu2* mutants defective in Ndc80c binding. Disrupting the Stu2:Ndc80c interaction led to diminished viability, aberrant chromosome biorientation, synthetic growth defects when combined with a *MAD3* deletion, and anaphase onset delays--all phenotypes consistent with defects in error correction. Chemically induced tethering of the mutant Stu2 to Ndc80c, rescued all the defects we examined, including viability and cell-cycle progression delays. Thus, the contributions of Stu2 to error correction *in vivo* depend on Ndc80 binding, not simply on its presence as a soluble factor or on its function as a microtubule polymerase. Our results do not rule out, however, that the required kinetochore localization of Stu2 couples its polymerase activity specifically with growth and shrinkage at the plus ends of kinetochore microtubules during metaphase. Its polymerase activity might therefore also be essential. Stu2 association with Bik1 and Spc72 presumably couple this activity with other stages of spindle assembly or with other roles for microtubules in a yeast cell.

A published study of Stu2 dynamics at kinetochores using fluorescence recovery after photobleaching showed relatively high turnover, and its authors suggested that Stu2 might not be a stably associated kinetochore factor (Aravamudhan et al., 2014). The bi-lobed clusters of Stu2 in metaphase cells (Goshima and Yanagida, 2000; He et al., 2001; Pearson et al., 2001) probably contain both kinetochore-bound and purely microtubule tip-associated Stu2 pools, however, making definitive interpretation difficult. Our current data show that regardless of its exchange rate, association of Stu2 with Ndc80c at the tetramer junction is necessary for its error correction function and that tethering Stu2 to the junction is sufficient to restore full activity of an association-defective mutant.

Conservation of the four-component organization of Ndc80c from yeast to mammals, despite no evidence for independent functions of either heterodimer, suggests a conserved role for the connection between them. The original plan when designing the dwarf construct was to provide a complex stiff enough to crystallize, while preserving the complete four-way junction. Several conserved features of the junction structure those crystals yielded, including exposed hydrophobic pockets and a skipped α-helical turn near the N-terminus of Spc24, indeed hinted that it might be a binding site for a conserved regulatory factor (Valverde et al., 2016). Stu2 is evidently at least one such factor. As now shown here, the Stu2 CTS adds a fifth helix to the four-helix bundle, without producing any conformational rearrangements, indicating that the role of the junction is simply to anchor Stu2 along the Ndc80c rod, rather than to receive an allosteric signal from it. That result does not rule out conformational switching by interactions between other parts of Stu2 thus anchored and other parts of the Ndc80c.

Bulky hydrophobic residues are present at key positions in the CTS of Stu2 orthologs from fungi to metazoans, and the entire CTS sequence has conserved features (Fig. 4E). This conservation suggests that the orthologs also bind Ndc80c. Published work provides evidence for interaction of the *S. pombe* ortholog with the Ndc80 internal loop (Hsu and Toda, 2011; Kakui et al., 2013). The C-terminal position of the Ndc80c attachment segment of Stu2 and the length of the presumably flexible linker between it and the dimerizing coiled-coil would allow more distal parts of the protein to interact with almost any other site along the Ndc80c rod, including the Ndc80 internal loop and the globular head (Miller et al., 2019), and probably even allow the TOG domains to contact the curled plus-ends of an attached microtubule. These additional interactions are likely to be critical contributors to the mechanism by which Stu2 responds to tension or its absence and synergizes with the error-correcting detachment activity of Ipl1.

## ACKNOWLEDGMENTS

We are grateful to Arshad Desai for providing antibodies, Sue Biggins, Leon Chan, Ajit Joglekar, and Frank Uhlmann for reagents, and to Sue Biggins for critical reading of the manuscript. This work was supported by a Damon Runyon-Dale F. Frey Award for Breakthrough Scientists (to M.P.M.). S.C.H. is an investigator of the Howard Hughes Medical Institute.

## MATERIALS AND METHODS

### Strain construction and microbial techniques

#### Yeast strains and plasmids

*Saccharomyces cerevisiae* strains used in this study, all derivatives of M3 (W303), are described in Table S1. Standard media and microbial techniques were used [48]. Yeast strains were constructed by standard genetic techniques. Construction of *pCUP1-GFP-LacI* and *ipl1-321* are described in (Biggins et al., 1999), *CEN3::lacO:TRP1* is described in (Shonn et al., 2003), *mad3Δ*, in (Pinsky et al., 2006), *DSN1-6His-3Flag*, in (Akiyoshi et al., 2010), *stu2-3V5-IAA7*, in (Miller et al., 2016). *TOR1-1, fpr1Δ*, and *MPS1-FRB:KanMX*, in (Aravamudhan et al., 2015; Haruki et al., 2008). *STU2-FRB:HisMX* and *NUF2-FKBP12:HisMX* were constructed by PCR-based methods (Longtine et al., 1998). Strains containing the previously described *pMET-CDC20* allele were provided by Frank Uhlmann. *pGPD1-TIR1* integration plasmids (pM76 for integration at *HIS3* or pM78 for integration at *TRP1)* were provided by Leon Chan. Construction of a 3HA-IAA7 tagging plasmid (pM69) was described previously (Miller et al., 2016). Construction of a *LEU2* integrating plasmid containing wild-type *pSTU2-STU2-3V5* (pM225) and *pSTU2-stu2(Δ855–888)-3V5* (pM267) are described in (Miller et al., 2016, 2019). STU2 variants were constructed by mutagenizing pM225 as described in (Liu and Naismith, 2008; Tseng et al., 2008). Primers used in the construction of the above plasmids are listed in Table S2, further details of plasmid construction including plasmid maps available upon request.

#### Auxin inducible degradation

The auxin inducible degron (AID) system was used essentially as described (Nishimura et al., 2009). Briefly, cells expressed C-terminal fusions of the protein of interest to an auxin responsive protein (IAA7) at the endogenous locus. Cells also expressed *TIR1*, which is required for auxin-induced degradation. 500 μM IAA (indole-3-acetic acid dissolved in DMSO; Sigma) was added to media to induce degradation of the AID-tagged protein. Auxin was added for 30 min prior to harvesting cells or as is indicated in figure legends.

#### Spotting assay

For the spotting assay, the desired strains were grown for 2 days on plates containing yeast extract peptone plus 2% glucose (YPD) medium. Cells were then resuspended to OD600 ~1.0 from which a serial 1:5 dilution series was made and spotted on YPD+DMSO, YPD+500 μM IAA (indole-3-acetic acid dissolved in DMSO) or plates containing 3.5-5.0 μg/mL benomyl or 0.05 μg/mL rapamycin as indicated. Plates were incubated at 23°C for 2-3 days unless otherwise noted.

#### Dissections

Diploid cells were sporulated, treated with 0.5 mg/ml zymolyase in 1 M sorbitol for 12 min, and tetrads were dissected on YPD plates using standard yeast microbial techniques. Plates were incubated at 23°C for 2 days.

#### FRB/FKBP re-tethering

For re-tethering in culture, exponentially growing cultures were treated with 500 μM auxin and 0.05 ug/mL or 55 nM rapamycin 30 min prior to harvesting. For spotting assays on plates, 0.05 ug/mL rapamycin was used.

#### Biorientation assay

In metaphase, sister kinetochores become bioriented and are transiently stretched apart by opposing microtubule pulling forces (Goshima and Yanagida, 2000; He et al., 2001; Pearson et al., 2001); the transient separation can be visualized by fluorescent marking of the centromere of a single chromosome (Straight et al., 1996). Biorientation was examined in metaphase arrested cells as judged by the fluorescently marked centromeres appearing as two distinct foci. Cells were grown in media lacking methionine (for *pMET-CDC20* containing strains). Exponentially growing cells were subsequently arrested in metaphase by the addition of 8mM methionine each hour for 3h. An aliquot of cells was fixed with 3.7% formaldehyde in 100mM phosphate buffer (pH 6.4) for 5 min. Cells were washed once with 100mM phosphate (pH 6.4), resuspended in 100mM phosphate, 1.2M sorbitol buffer (pH 7.5) and permeabilized with 1% Triton X-100 stained with 1 μg/ml DAPI (4’, 6-diamidino-2-phenylindole; Molecular Probes).

#### Cell cycle progression assay

Cells were grown in YPD medium. Exponentially growing MATa cells also carrying a tandem array of lacO sequences integrated proximal to CEN3 (Shonn et al., 2003) and a LacI-GFP fusion (Biggins et al., 1999; Straight et al., 1996) were arrested in G1 with 1μg/ml α-factor. When arrest was complete, cells were released into medium lacking α-factor pheromone and containing 500μM IAA and 0.05 ug/mL rapamycin at 23°C. ~75 min after G1 release, 1μg/ml α-factor was added to prevent a second cell division. Samples were taken every 15min after G1 release to determine cell cycle progression (via nuclear morphology of DAPI stained nuclei).

### Protein biochemistry

#### Purification of native kinetochore particles

Native kinetochore particles were purified from asynchronously growing *S. cerevisiae* cells as described below. Dsn1-6His-3Flag was immunoprecipitated with anti-Flag (essentially as described in (Akiyoshi et al., 2010). Cells were grown in yeast peptone dextrose rich (YPD) medium. For strains containing Stu2-AID, cells were treated with 500 μM auxin 30 min prior to harvesting. Protein lysates were prepared by mechanically disrupted in the presence of lysis buffer using glass beads and a beadbeater (Biospec Products). Lysed cells were resuspended in buffer H (BH) (25 mM HEPES pH 8.0, 2 mM MgCl2, 0.1 mM EDTA, 0.5 mM EGTA, 0.1% NP-40, 15% glycerol with 150 mM KCl containing protease inhibitors (at 20 μg mL-1 final concentration for each of leupeptin, pepstatin A, chymostatin and 200 μM phenylmethylsulfonyl fluoride) and phosphatase inhibitors (0.1 mM Na-orthovanadate, 0.2 μM microcystin, 2 mM β-glycerophosphate, 1 mM Na pyrophosphate,5 mM NaF) followed by centrifugation at 16,100 g for 30 min at 4°C to clarify lysate. Dynabeads conjugated with anti-Flag antibodies were incubated with extract for 3 h with constant rotation, followed by three washes with BH containing protease inhibitors, phosphatase inhibitors, 2 mM dithiothreitol (DTT) and 150 mM KCl. Beads were further washed twice with BH containing 150 mM KCl and protease inhibitors. Associated proteins were eluted from the beads by boiling in 2x SDS sample buffer.

#### Immunoblot analysis

For immunoblot analysis, cell lysates were prepared as described above or by pulverizing cells with glass beads in sodium dodecyl sulfate (SDS) buffer using a bead-beater (Biospec Products). Standard procedures for sodium dodecyl sulfate-polyacrylamide gel electrophoresis (SDS-PAGE) and immunoblotting were followed as described in (Burnette, 1981; Towbin et al., 1992). A nitrocellulose membrane (Bio-Rad) was used to transfer proteins from polyacrylamide gels. Commercial antibodies used for immunoblotting were as follows: α-Flag, M2 (Sigma-Aldrich) 1:3,000; α-V5 (Invitrogen) 1:5,000. Antibodies to Ndc80 were a kind gift from Arshad Desai and used at anti-Ndc80 (OD4) 1:10,000. The secondary antibodies used were a sheep anti-mouse antibody conjugated to horseradish peroxidase (HRP) (GE Biosciences) at a 1:10,000 dilution or a donkey anti-rabbit antibody conjugated to HRP (GE Biosciences) at a 1:10,000 dilution. Antibodies were detected using the SuperSignal West Dura Chemiluminescent Substrate (Thermo Scientific).

#### Recombinant protein expression and purification

The constructs used to express the dwarf Ndc80c are identical to those used previously (Valverde et al., 2016). Briefly, shortened versions of Ndc80/Nuf2, and Spc24/Spc25 were cloned into the orthogonal expression vectors pETduet1 and pRSFduet, respectively. Proteins were co-expressed in Rosetta 2(DE3)pLysS Ecoli (Novagen). Cells were grown at 37°C in 2XYT media to an OD600 of 0.8, induced with 200 μM IPTG, and incubated overnight with shaking at 225 RPM at 18°C. Cell pellets from 6L of culture were resuspended in 150 mL of buffer containing 50 mM Tris pH 8.0, 250 mM NaCl, 10 mM imidazole pH 8.0, 5 mM β-mercaptoethanol, 1 mM PMSF, 1 μg/mL pepstatin, 1 μg/mL aprotinin, 1 μg/mL leupeptin, 83 μg/mL lysozyme, 30 μg/mL Dnase I. Cells were lysed by sonication, and the lysate was clarified by centrifugation and applied to Ni-NTA agarose equilibrated with buffer containing 20 mM Tris pH 8.0, 100 mM NaCl, 10 mM imidazole pH 8.0, and 5 mM β-mercaptoethanol (equilibration buffer). Bound material was washed with equilibration buffer containing 20 mM imidazole and 500 mM NaCl and then with equilibration buffer. Proteins were eluted with equilibration buffer containing 400 mM imidazole pH 8.0 and 50 mM NaCl. Eluates were treated overnight with TEV protease to remove the 6-His tag from the N-terminus of Ndc80 (except protein immobilized for pulldown experiments) and subjected to anion exchange chromatography using a Hi-trap Q HP (Cytiva), followed by size-exclusion chromatography using a Hi-load Superdex 200 16/60 column (Cytiva) equilibrated with 10 mM HEPES pH 7.0, 100 mM NaCl, and 2 mM tris(2-carboxyethyl)phosphine (TCEP).

Full length Stu2 was expressed in SF9 cells using the Bac-To-Bac system. Following infection, protein expression was allowed to proceed for 72 hours at 27°C. Cells were resuspended in 50 mM Tris pH 7.2, 500 mM NaCl, 10 mM imidazole pH 7.2, 5 mM β-mercaptoethanol, 0.1% TWEEN-20, 0.1% IPGAL, 1 mM PMSF, 1 μg/mL pepstatin, 1 μg/mL aprotinin, 1 μg/mL leupeptin, and 30 μg/mL dnase I. Cells were lysed using a dounce homogenizer. The lysate was clarified by centrifugation and applied to Ni-NTA agarose equilibrated with 20 mM HEPES pH 7.2, 100 mM NaCl, 10 mM imidazole pH 7.2, and 5 mM β-mercaptoethanol. Resin was washed with buffer containing 500 mM NaCl and 20 mM imidazole, washed again with equilibration buffer and eluted with buffer containing 400 mM Imidazole and 50 mM NaCl. Eluates were then subjected to cation exchange chromatography using a 5 mL Hi-Trap SP HP column (Cytiva). Fractions containing full-length Stu2 were applied to a Superose 6 10/300 column (Cytiva) equilibrated with 10 mM tris pH 7.5, 150 mM Nacl and 2 mM TCEP.

The C-terminal fragments of Stu2 used in the pulldown experiments were cloned into a pLIC vectors with a TEV-cleavable N-terminal 6-His tag and transformed into Bl-21 AI cells (Invitrogen). Cells were grown to an OD_600_ of 0.6 in TB media with 0.1% glucose and were induced by addition of IPTG and arabinose to 200 μM and 0.2%, respectively, incubated at 37°C for 4 hours to consume the glucose and then at 18° overnight. Cells were harvested by centrifugation, resuspended in 50 mM Tris pH 7.2, 250 mM NaCl, 10 mM Imidazole pH 8.0, 5 mM β-mercaptoethanol, 1 mM PMSF, 1 μg/mL pepstatin, 1 μg/mL aprotinin, 1 μg/mL leupeptin, 83 μg/mL lysozyme, 30 μg/mL Dnase, and lysed by sonication. The lysates were clarified by centrifugation and applied to Ni-NTA agarose equilibrated with 20 mM HEPES pH 7.2, 100 mM NaCl, 10 mM imidazole pH 7.2, and 5 mM β-mercaptoethanol. Eluates were treated with TEV protease overnight then subjected to cation exchange chromatography on a 5mL Hi-Trap SP HP column (Cytiva). Fractions containing Stu2 were then applied to a Superdex200 10/300 column (Cytiva) equilibrated with 10 mM HEPES pH 7.2, 150 mM NaCl, and 2 mM TCEP.

#### Peptide

A 33-mer C-terminal Stu2 peptide, with a selenomethionine in place of the natural methionine at position 876 (EESYKRAAAVTSTLKARIEK(Mse)KAKSRREGTTRT), was synthesized at the Tufts University Core facility. The lyophilized peptide was resuspended to a final concentration of 10 mg/mL in a buffer containing 10 mM TRIS pH 7.0, 100 mM NaCl and 2 mM TCEP.

### Crystallization, diffraction data collection, and structure determination

Dwarf Ndc80c was concentrated to 15 mg/mL in an Amicon centrifugal concentrator in buffer containing 10 mM HEPES pH 7.0, 100 mM NaCl, and 2 mM TCEP. The Se-Met Stu2 peptide was diluted to 2 mg/mL in the same buffer. The protein was crystallized using hanging-drop vapor diffusion with a well solution containing 1.3M ammonium sulfate and 0.1 mM HEPES pH 7.3. The hanging drops contained 1 μL each of the Ndc80c, peptide and well solutions, resulting in a ~3-fold molar excess of peptide over dwarf Ndc80c. Crystals were cryoprotected in well solution supplemented with 25% glucose and flash-frozen by plunging into liquid N_2_. The complex crystallized in the C222_1_ space group (a = 190.89 Å, b=183.3 Å, c = 124.32 Å). Data to a minimum Bragg spacing of 2.7 Å were recorded on the NE-CAT beamline 24-ID-C at the Advanced Photon Source, indexed and integrated using HKL2000, and scaled and merged using Scalepack as implemented in HKL2000 (Otwinowski and Minor, 1997) (Table S3). The structure was determined by molecular replacement in Phenix (Adams et al., 2010), using the dwarf Ndc80c (PDB 5TCS) as a search model, yielding clear density for the Stu2 peptide. Model building was carried out in Coot (Emsley et al., 2010) and refinement, in Phenix (Adams et al., 2010) (Table S3). We used the anomalous signal from a Se-methionine residue to calculate an anomalous difference map, confirming that the Se atom was at a position consistent with the chosen register. Coordinates and diffraction data have been deposited in the protein data bank, PDB ID: 7KDF

### Pulldown experiments

Dwarf Ndc80c with a 6-His affinity tag on the N-terminus of Ndc80 was immobilized to saturation on Co-NTA agarose by incubation with gentle agitation at 4°C. Beads were pelleted by centrifugation, washed three times in 20 mM HEPES pH 7.5, 200 mM NaCl, 2 mM imidazole pH 7.2, 2 mM β-mercaptoethanol, and 0.1% TWEEN-20 (wash buffer), and incubated with the relevant Stu2 construct for 30 minutes. Beads were again pelleted, washed three times with wash buffer, pelleted a final time, and the wash buffer aspirated from the tube. To elute bound proteins, 50 μL of wash buffer supplemented with 500 mM imidazole pH 7.5 was added to ~25 μL of beads. After an additional spin, the eluted proteins were visualized by SDS-PAGE.

## SUPPLEMENTAL FIGURE LEGENDS

**Supplement to Figure 3.**
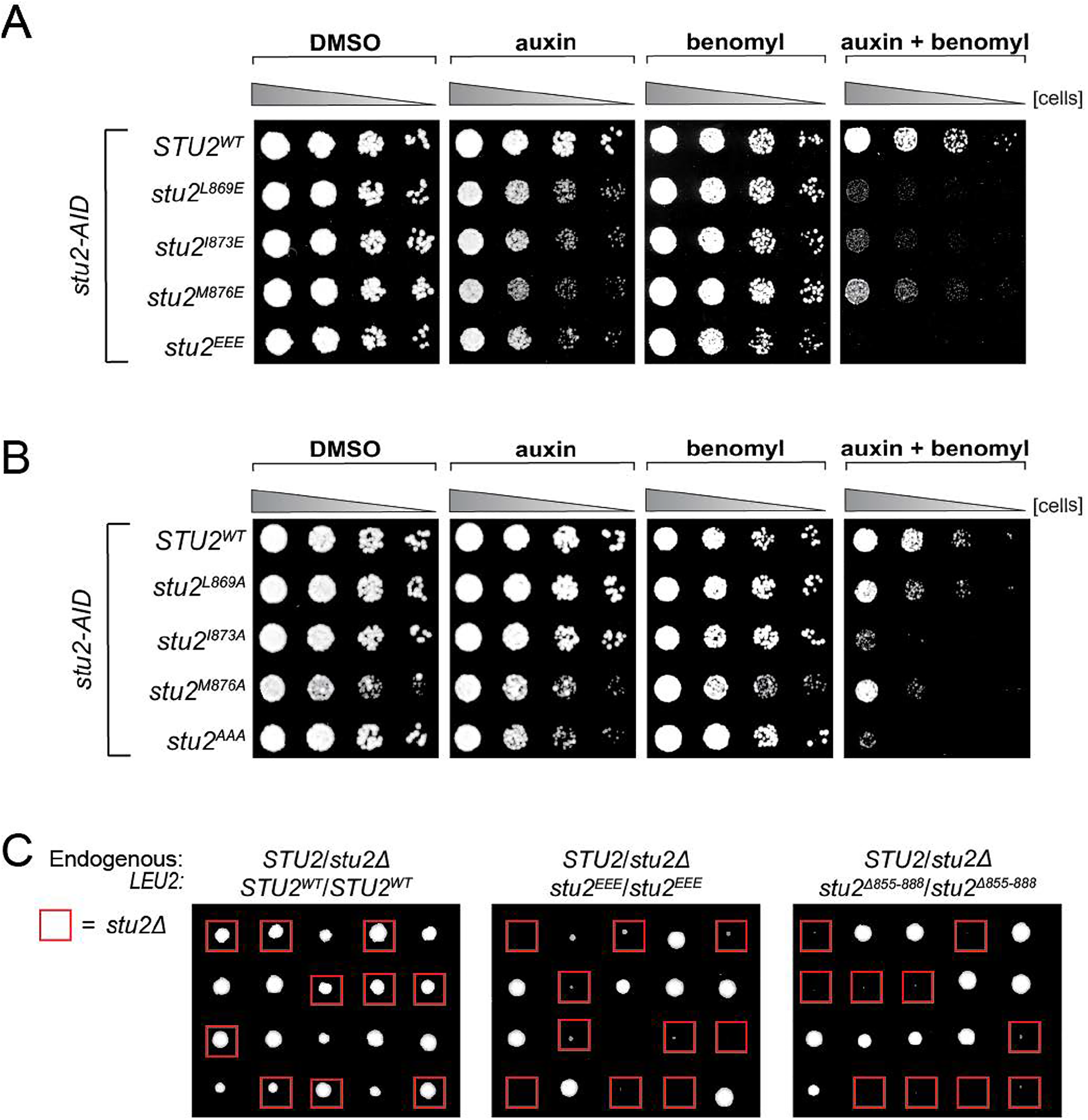
**A.** *stu2-AID* cells expressing various *STU2-3V5* alleles from an ectopic locus (*STU2^WT^*, M622; *stu2^L869E^*, M1443; *stu2^I873E^*, M1442; *stu2^M876E^*, M1441; *stu2^EEE^*, M1444) were serially diluted (5-fold) and spotted on plates containing DMSO, 500 μM auxin, 5 μg/mL benomyl or 500 μM auxin + 5 μg/mL benomyl. **B.** *stu2-AID* cells expressing various *STU2-3V5* alleles from an ectopic locus (*STU2^WT^*, M622; *stu2^L869A^*, M1576; *stu2^I873A^*, M1575; *stu2^M876A^*, M1574; *stu2^AAA^*, M1525) were serially diluted (5-fold) and spotted on plates containing DMSO, 500 μM auxin, 5 μg/mL benomyl, or 500 μM auxin + 5 μg/mL benomyl. **C.** Diploid cells heterozygous for *STU2/stu2Δ::HIS3* at the endogenous *STU2* locus and homozygous for different *STU2* alleles at the *LEU2* locus (*STU2^WT^/STU2^WT^*, M1718; *stu2^EEE^/stu2^EEE^*, M1714; *stu2^Δ855-888^/stu2^Δ855-888^*, M1716) were sporulated and dissected on YPD plates. Spores scored positively for HIS3 are boxed in red. Spores harboring both deletion of endogenous *STU2* and either *stu2^EEE^* or *stu2^Δ855-888^* have a viability defect. The *stu2^EEE^* strain shows a dominant meiotic phenotype in this assay as well, possibly resulting from aberrant meiotic chromosome segregation in *STU2/stu2Δ:HIS3 stu2^EEE^/stu2^EEE^* diploid cells.

**Supplement to Figure 4.**
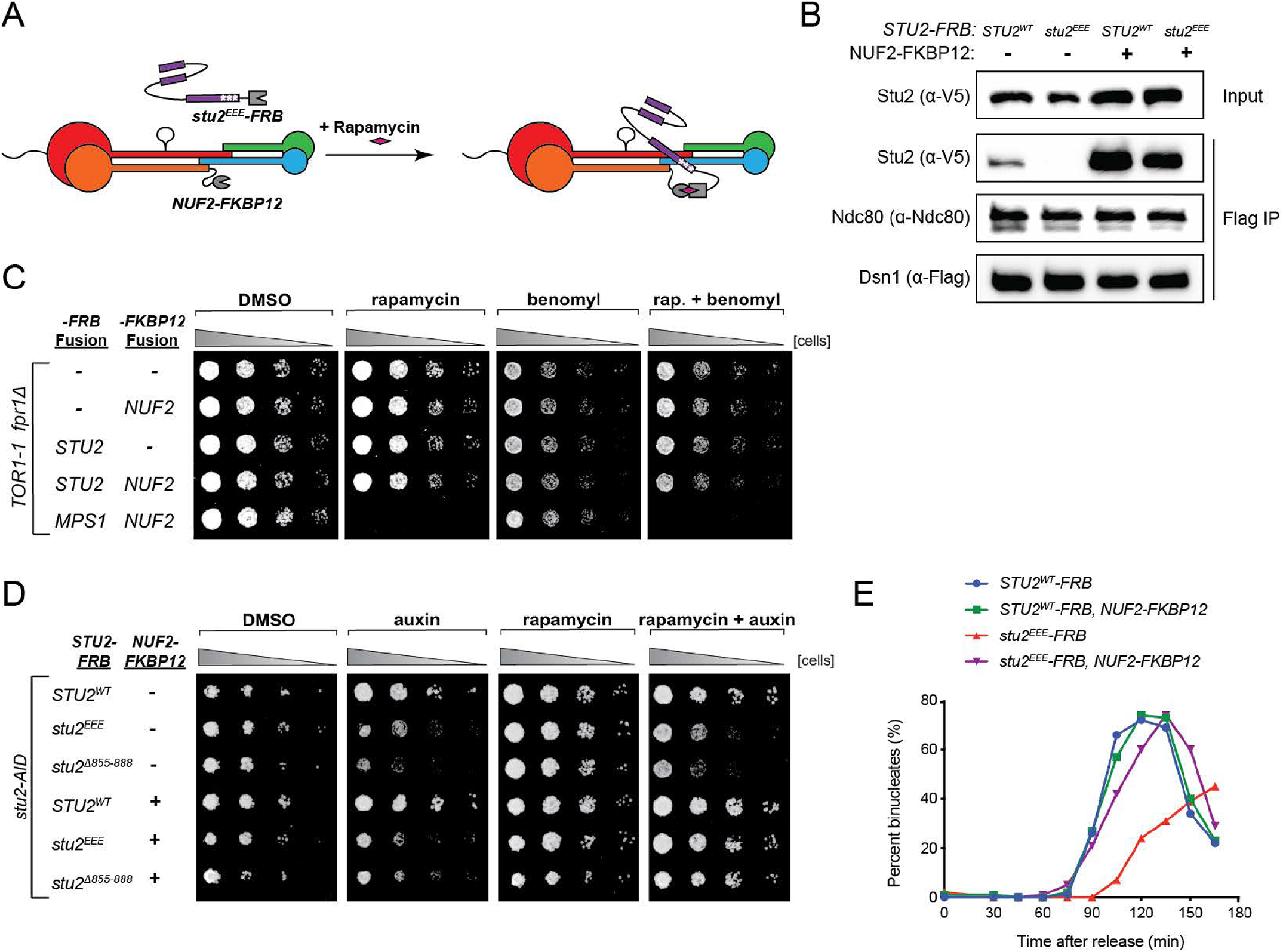
**A.** Schematic illustrating re-tethering of *stu2^EEE^* to Ndc80c. In the presence of rapamycin, *stu2^EEE^-FRB* binds to *NUF2-FKBP12*, restoring *stu2^EEE^*-Ndc80c interaction. **B.** Reproduction of western blot from Figure 3D also showing immunoblot signal for Dsn1-6His-3Flag. **C.** *fpr1Δ TOR1-1* cells (M1375), and *fpr1Δ TOR1-1* cells expressing-*FRB* and/or -*FKBP12* fusion alleles at the endogenous fusion-target locus (*NUF2-FKBP12*, M1422; *STU2-FRB*, M1387; *STU2-FRB NUF2-FKBP12*, M1428; *MPS1-FRB NUF2-FKBP12*, M1461) were serially diluted (5-fold) and spotted on plates containing DMSO, 50 ng/mL rapamycin, 3.5 ug/mL benomyl, or 3.5 ug/mL benomyl + 50 ng/mL rapamycin. Note: tethering Stu2-FRB to Nuf2-FKBP12 does not affect cell viability, whereas tethering Mps1-FRB results in lethality, as previously reported, (Aravamudhan et al., 2015) **D.** *stu2-AID fpr1Δ TOR1-1* cells expressing *STU2-FRB* alleles at an ectopic locus with and without *NUF2-FKBP12* (*STU2^WT^-FRB*, M1513; *stu2^EEE^-FRB*, M1515; *stu2^Δ855-888^-FRB*, M1587; *STU2^WT^-FRB NUF2-FKBP12*, M1505; *stu2^EEE^-FRB NUF2-FKBP12*, M1507; *stu2^Δ855-888^-FRB NUF2-FKBP12*, M1554) were serially diluted (5-fold) and spotted on YPD plates containing DMSO, 50 ng/mL rapamycin, 500 μM auxin, or 500 μM auxin + 50 ng/mL rapamycin. Tethering *stu2^Δ855-888^-FRB* to Ndc80c restores viability to the same degree as does tethering *stu2^EEE^-FRB*. Thus the essential function of the Stu2 CTS is kinetochore binding. These data also imply that the kinetochore function of Stu2 does not require its association with other factors that bind the CTS. **E.** Replicate cell cycle delay experiment as described in Figure 4D

**Table S1.**
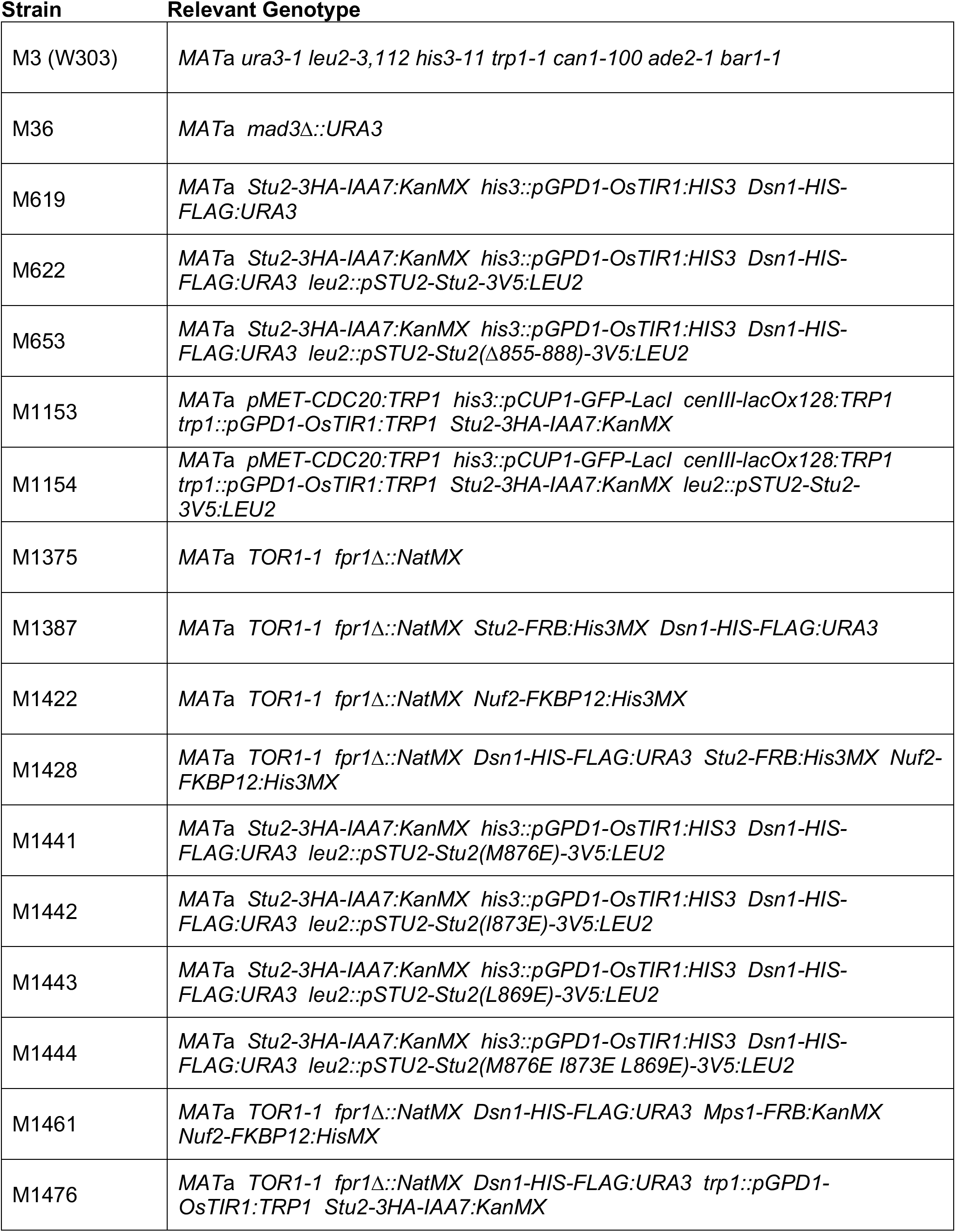
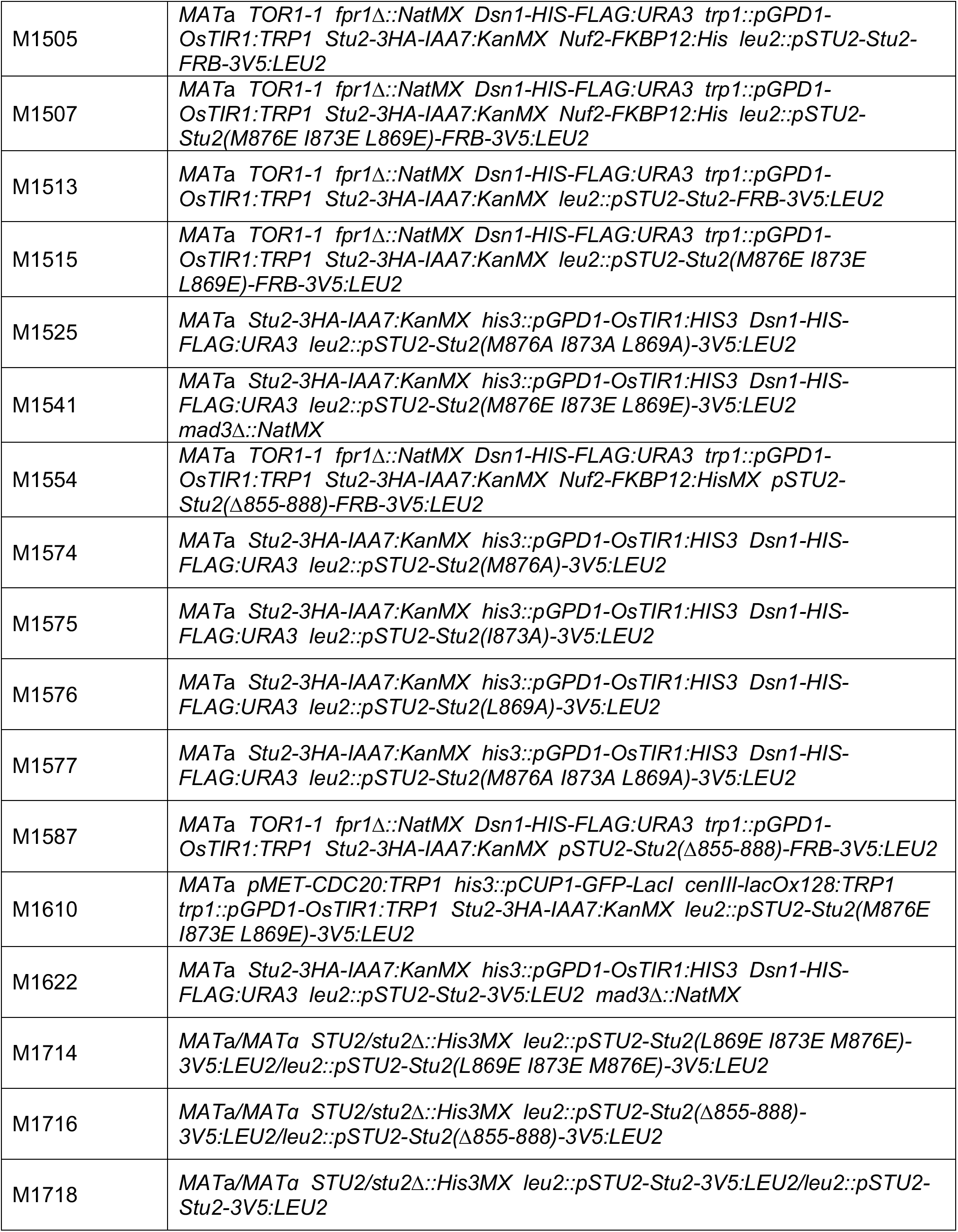
Strains used in this study. All strains are derivatives of M3 (W303)

**Table S2.**
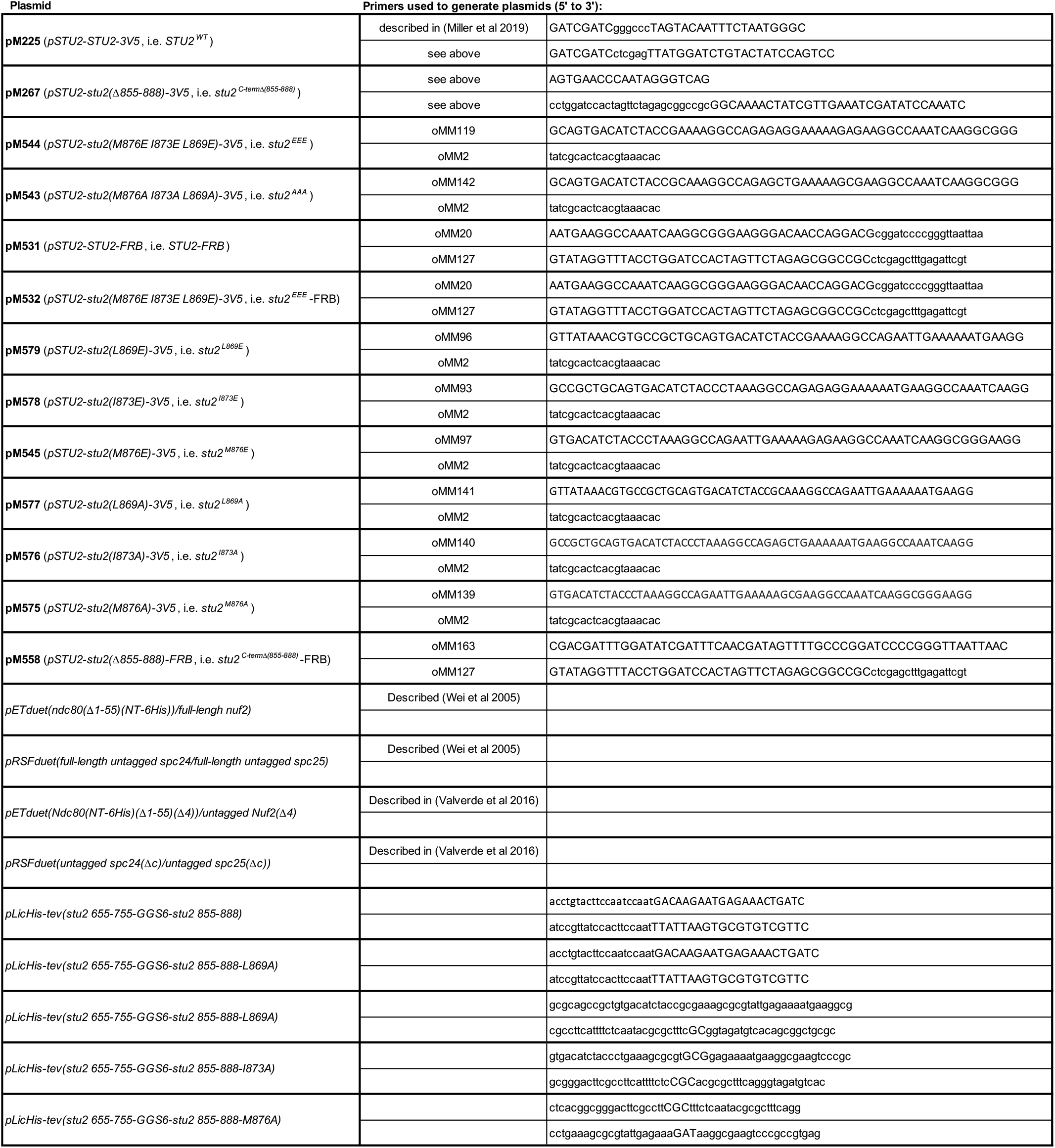
Plasmids and Primers used in this study. All pM plasmids are derivatives of pM225.

**Table S3.** Data collection and refinement statistics

|  | Stu2 CTS bound to dwarf Ndc80c* |
| --- | --- |
| Wavelength | 0.97918 Å |
| Resolution range | 42.72 - 2.72 (2.817 - 2.72) |
| Space group | C 2 2 21 |
| Unit cell | 170.886 183.175 124.317 90 90 90 |
| Total reflections | 697683 (62919) |
| Unique reflections | 52106 (4719) |
| Multiplicity | 13.4 (13.3) |
| Completeness (%) | 99.03 (91.15) |
| Mean I/sigma(I) | 18.40 (1.78) |
| Wilson B-factor | 63.68 |
| R-merge | 0.1071 (0.9043) |
| R-meas | 0.1114 (0.9395) |
| R-pim | 0.03047 (0.2529) |
| CC1/2 | 0.999 (0.829) |
| CC* | 1 (0.952) |
| Reflections used in refinement | 52103 (4719) |
| Reflections used for R-free | 2013 (194) |
| R-work | 0.1976 (0.2789) |
| R-free | 0.2416 (0.2910) |
| CC(work) | 0.962 (0.822) |
| CC(free) | 0.960 (0.757) |
| Number of non-hydrogen atoms | 5994 |
| macromolecules | 5869 |
| ligands | 30 |
| solvent | 95 |
| RMS(bonds) | 0.005 |
| RMS(angles) | 0.74 |
| Ramachandran favored (%) | 97.57 |
| Ramachandran allowed (%) | 2.43 |
| Ramachandran outliers (%) | 0.00 |
| Rotamer outliers (%) | 0.15 |
| Clashscore | 6.46 |
| Average B-factor | 88.87 |
| macromolecules | 89.23 |
| ligands | 98.66 |
| solvent | 63.61 |
| Number of TLS groups | 1 |

\*Statistics for the highest-resolution shell are shown in parentheses.

